# Impaired perception of isoluminant contrast modulation stimuli: Evidence for a magnocellular pathway mechanism

**DOI:** 10.1101/2025.06.04.657905

**Authors:** Ana L. Ramirez, Anna Shakhgildian, Ari Rosenberg, Curtis L. Baker

## Abstract

Contrast modulation (CM) stimuli have been previously used to reveal nonlinear contributions of Y-like retinal ganglion cells (RGCs) such as parasol cells to cortical responses and perception. To test whether CMs are selectively processed within the magnocellular pathway, we assessed envelope motion discrimination and detection for achromatic (yellow-black) and chromatic (red-green) CMs in the presence of luminance masking noise to disrupt luminance-based mechanisms of motion processing. Compared to achromatic CMs, perception of chromatic CMs was more sensitive to luminance masking noise, suggesting that CM envelope motion perception relied predominantly on luminance signals. Specifically, envelope motion discrimination performance was better maintained for achromatic CMs than chromatic CMs, even at high masking noise levels. Notably, luminance masking noise greatly impaired envelope direction discrimination for chromatic CMs but had minimal impact on their detection, suggesting that chromatic aberrations may enhance envelope motion perception for chromatic CMs by introducing luminance signals. These findings collectively emphasize that CM stimuli selectively activate Y-like/parasol RGCs of the retino-geniculate magnocellular pathway, underscoring their potential to specifically target this pathway. This might have clinical advantages for early diagnosis in disorders such as glaucoma or dyslexia, where magnocellular pathway dysfunction is predominant.

## Introduction

The retina is the nexus between light and perception, providing the basis of visual experience. In primates, two major classes of retinal ganglion cells (RGCs) are midget cells and parasol cells (Rodieck, 1979; Dacey & Lee, 1994). Midget cells belong to the parvocellular pathway, have smaller receptive fields, and lower contrast sensitivity. They compose about 80% of the population of the central retina where high spatial resolution is crucial (Leventhal et al., 1981; Perry et al., 1984). Lesions to the parvocellular geniculocortical pathway decrease contrast sensitivity to visual stimuli with high spatial frequencies (SFs) and low temporal frequencies (TFs), compromise visual acuity, and cause a severe reduction in color vision (Merigan et al 1991a,b). Parasol cells belong to the magnocellular pathway, have larger receptive fields and higher contrast sensitivity, and respond to higher TFs (Kaplan and Shapley, 1982; Leventhal et al., 1981; Perry et al., 1984). Lesions to the magnocellular pathway decrease contrast sensitivity for visual stimuli at low SFs and high TFs (Merigan et al 1991a).

Parasol cells in the macaque retina are now recognized to have nonlinear properties like those of Y-type cells in the cat (Crook et al., 2008a). They exhibit a first harmonic (F1) response to low SF drifting sinewave gratings, which indicates a linear response at the TF of the stimulus. However, they also manifest a second harmonic (F2) response to high SF contrast-reversing gratings, which remains consistent across spatial phases. The F2 responses come from the activation of small nonlinear subunits, originating from rectified cone bipolar cell inputs (Demb et al., 1999). This phase invariant F2 response is characteristic of the nonlinear response exhibited by Y cells (Hochstein and Shapley, 1976; Kaplan and Shapley 1982; Shapley and Perry, 1986). Consequently, here we consider parasol cells to be “Y-like”.

Contrast modulation (CM) stimuli are comprised of a high SF carrier pattern (e.g., a sinewave grating) whose contrast is modulated by a low SF sinewave envelope. They are considered “second order” stimuli since they are characterized by spatial variations in texture or local contrast as opposed to luminance (Cavanagh & Mather, 1989). Neurophysiology studies have demonstrated that CM stimuli activate early visual cortex neurons in cats (Zhou and Baker, 1994; Mareschal and Baker, 1998) and macaque monkeys (Li et al., 2014). In cats, LGN Y cells respond to CM stimuli with similar selectivity for the carrier orientation and SF as cortical neurons. This similarity suggests the likely involvement of a neural mechanism driven by nonlinear subunits of subcortical Y cells (Rosenberg et al., 2010; Rosenberg and Issa, 2011).

Additionally, both LGN Y cells (Rosenberg et al., 2010) and cortical area 18 neurons (Rosenberg and Issa, 2011; Gharat and Baker, 2012) respond to CM stimuli with surprisingly high carrier TFs, suggesting that these responses are unlikely to originate from cortical feedback mechanisms. These findings collectively support the idea that nonlinear processing of CM stimuli in the cortex fundamentally depends on input from retinal Y-like cells (Rosenberg and Talebi, 2009).

Recent psychophysical work used CM stimuli to reveal nonlinear contributions of Y-like cells to human visual perception (Ramirez et al., 2022). In those experiments, motion direction discrimination of luminance modulation (LM) gratings and the envelopes of CM stimuli at different carrier SFs and TFs was measured. Drifting LM stimuli yielded optimal performance at combinations of low TFs and high SFs, or vice versa. The strong interaction of the stimulus parameters was consistent with previous results for sinewave grating psychophysics (Robson, 1966; Watson & Ahumada, 2016). In contrast, performance for CM stimuli remained good at combinations of high carrier SFs and high carrier TFs, without an apparent interaction. The CM results also revealed a bandpass dependence on carrier SF consistent with findings on CM-responsive neurons in cats (Zhou & Baker, 1996; Mareschal & Baker, 1999) and monkeys (Li et al., 2014). In addition, Ramirez et al. (2022) examined performance for the same stimuli and task while varying stimulus eccentricity. The LM performance declined systematically with eccentricity, as conventionally expected, whereas CM performance remained more stable. These results may reflect previous neurophysiology findings that, in primate retina, the center F1 (linear) receptive field size of parasol cells increases substantially with eccentricity whereas the center F2 (nonlinear) receptive field size varies minimally with eccentricity (Crook et al., 2008a). Overall, the alignment between psychophysical observations and neurophysiological evidence suggests that perception of CM stimuli reflects nonlinear processing by parasol cells. As such, CM stimuli may offer a selective tool for assessing the function of parasol cells, which could potentially allow for earlier detection of pathologies associated with the magnocellular pathway, such as glaucoma (Quigley et al., 1987, 1988; Glovinsky et al., 1991; Shou et al., 2003; Zhang et al., 2016) and developmental dyslexia (Livingstone et al., 1991; Laprevotte et al., 2021).

Because parasol cells are largely achromatic (Leventhal et al., 1981; Perry et al., 1984) in their selectivity, magnocellular-mediated mechanisms of motion processing are expected to respond poorly to isoluminant chromatic stimuli. Motion direction discrimination for stimuli modulated by isoluminant color is a subject of ongoing debate. Studies have yielded diverse results, including perceived slower motion and momentary disappearance of motion for isoluminant red-green stimuli compared to luminance-based counterparts (Cavanaugh et al., 1984; Livingstone & Hubel, 1988; Mullen & Boulton 1992a, 1992b), but good apparent motion in some circumstances (Cropper & Derrington, 1996; Dobkins & Albright, 1993), as well as a motion aftereffect (Mullen & Baker, 1985). These discrepant results might reflect differences in the inadvertent activation of luminance-based mechanisms by chromatic aberrations arising from optical imperfections (Flitcroft, 1989; Bradley et al., 1992) or from differing temporal delays in the processing of long-(L) and middle-wavelength (M) cone signals (Bradley et al., 1992; Faubert et al., 2000; Mullen et al., 2003), both of which can create spurious luminance signals (see Discussion). In such cases, superimposing luminance noise on chromatic stimuli can be used to mask the contributions of luminance mechanisms to motion direction discrimination (Gegenfurtner & Kiper, 1992; Losada & Mullen, 1995), but not necessarily detection (Mullen et al., 2003; Yoshizawa et al., 2003).

Here we test the hypothesis that magnocellular processing mediates perception of CM stimuli, by comparing psychophysical measurements of motion direction discrimination for achromatic and isoluminant chromatic CM stimuli. To assess whether the activation of luminance-based mechanisms contributes to the perception of chromatic CM stimuli, we incorporate varying levels of luminance masking noise. Our findings suggest that discriminating the direction of chromatic CM stimuli relies on luminance-based mechanisms that are effectively masked by luminance noise. This result supports the idea that human visual perception of CM motion stimuli depends on magnocellular processing, consistent with neurophysiological findings from cats and monkeys.

## Methods

### Subjects

Six subjects (aged 19–37 years; 5 females, 1 male) participated. Four were naive to the aims of the study and two were authors. All subjects had a monocular, best-corrected visual acuity of at least 20/30 and reported no history of ophthalmological diseases or surgeries. All procedures were approved by the Research Ethics Board of the Research Institute of McGill University Health Center and were performed according to relevant guidelines and regulations, including the tenets of the Declaration of Helsinki. All subjects gave written informed consent to participate.

### Apparatus

Visual stimuli were produced on a PC (Intel i5, 6-core; 32 GB; NVIDIA GT 1030) running Linux (Kubuntu 20.04) with custom software written in MATLAB (MathWorks, Inc.) using the Psychophysics Toolbox, version 3.0.10 (Brainard, 1997; Kleiner, 2007; Pelli, 1997). The stimuli were presented on a cathode-ray tube (CRT) monitor (Iiyama, 39.5 × 29.5 cm, 120 Hz, 1024 × 768 pixels) at a viewing distance of 221 cm. We measured the gamma nonlinearity separately for the red, green, and blue guns of the CRT with a photometer (United Detector Technology S370), applied inverse lookup table correction, and verified linearization.

### Stimuli

The stimuli were presented at the center of the CRT monitor screen within a 3.5° cosine-tapered circular window, with a fixation target in the upper right quadrant such that the stimuli were located at 4.3° of retinal eccentricity (Figure 1). Subjects viewed the stimuli monocularly with the right eye. The left eye was occluded.

**Figure 1.**
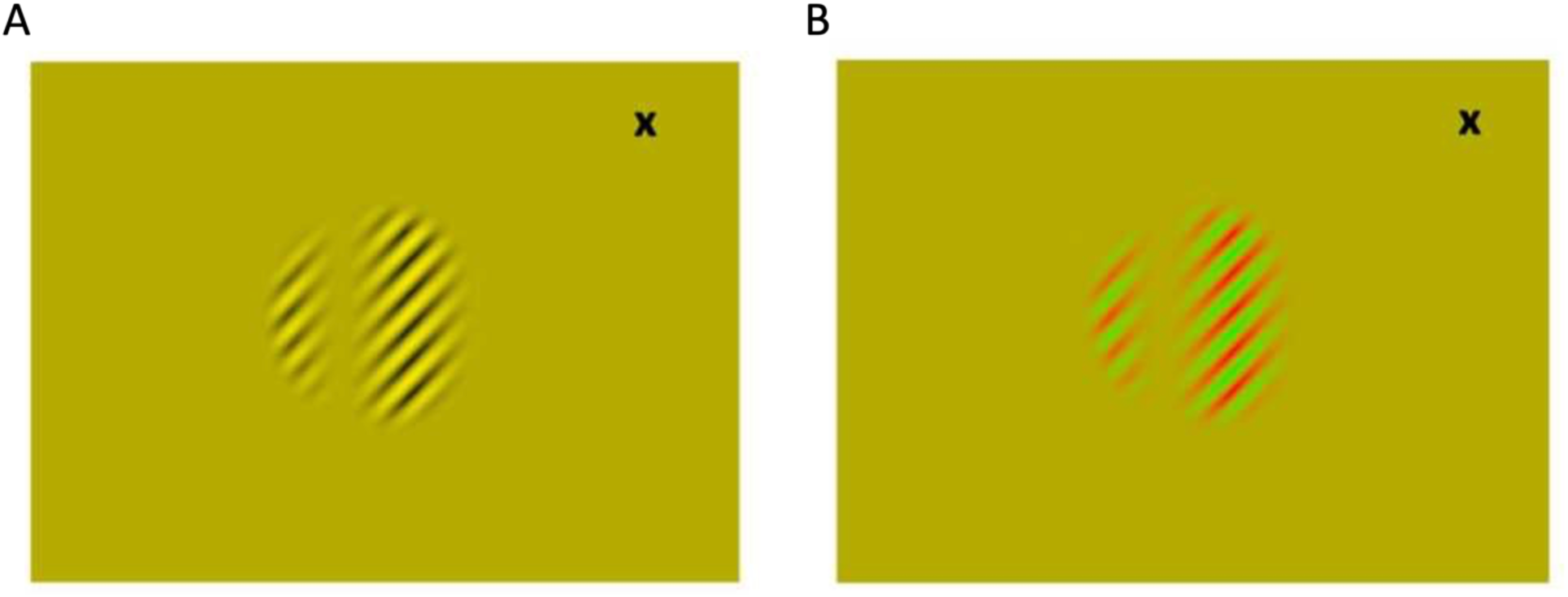
Example contrast modulation (CM) stimuli on a yellow background, with an eccentric fixation target (“x”) in the upper right quadrant. A) Achromatic yellow-black (Y-B) CM stimulus with a right-oblique carrier (high spatial frequency, SF) and vertical envelope (low SF). B) Same as A, but for a chromatic red-green (R-G) CM stimulus. For purposes of illustration, the SFs shown here are not at the same scale as used in the experiments.

A CM grating was defined by the contrast modulation of a high SF carrier grating by a low SF envelope grating:

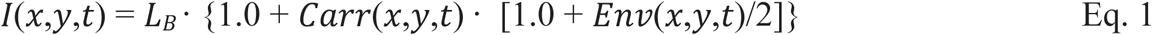

where *I*(*x*,*y*,*t*) was the intensity of a pixel at spatial location (*x*,*y*) at time *t*, *L*_*B*_ was the background (and mean) intensity, and *Carr*(*x*,*y*,*t*) was the carrier pattern, which was a contrast-reversing grating:

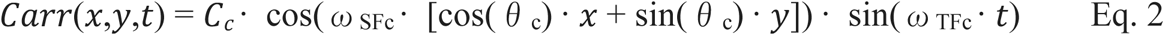

where *C*_*c*_ was the carrier contrast, ω_SFc_ was the carrier SF, θ_c_ was the carrier orientation, and ω_TFc_ was the carrier TF. The carrier orientation was always right-oblique, as in Figure 1.

The envelope, *Env*(*x*,*y*,*t*), was a drifting grating:

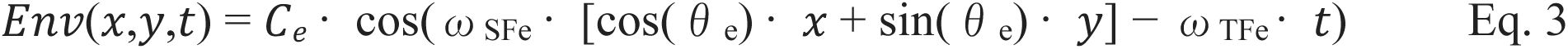

where *C*_*e*_ was the envelope contrast, ω_SFe_ was the envelope SF, θ_e_ was the envelope orientation, and ω_TFe_ was the envelope TF. The envelope orientation was always vertical, as in Figure 1, so its motion was either leftwards or rightwards. The orientation, SF, and TF parameters were defined independently for the carrier and the envelope.

We presented both achromatic yellow-black (Y-B) CM stimuli as well as isoluminant chromatic red-green (R-G) CM stimuli against a uniform yellow background (Figure 1). By “achromatic”, we mean that the stimuli were modulated in luminance but not in chromaticity (Baker et al., 1998; Metha & Mullen, 1997; Shooner & Mullen, 2020). The achromatic (Y-B) (Figure 1A) and chromatic (R-G) (Figure 1B) CM stimuli were produced with gamma-corrected color lookup tables for the red and green guns of the CRT that were acting in concert (Y-B) or opposition (R-G), with the blue gun disabled (i.e., all blue lookup table values were set to zero).

Additionally, spatially 1-D, achromatic (Y-B) masking noise was presented within the cosine-tapered circular window where the CM stimuli were displayed. The masking noise was vertically oriented, to be maximally effective for the vertical envelopes of the CM stimuli. The noise was generated with uniformly distributed random values, with fresh random samples used for each frame of a given trial and for each trial. A1-D spatial low-pass filter (Butterworth, order = 14) was applied to each frame of the broadband noise, with a cutoff of 2 cycles per degree (cpd). The RMS contrast (standard deviation divided by mean) of each frame was then set to the desired value.

The CM stimuli and masking noise were presented on alternate frames of the 120 Hz CRT monitor; that is, each was updated at 60 Hz. As necessary, the color lookup tables were also changed with each frame to produce either achromatic (Y-B) or chromatic (R-G) CM stimuli and achromatic (Y-B) masking noise. Consequently, the effective (time-averaged) contrasts of both the CM stimuli and masking noise were one-half the specified values.

### Experimental procedures

We verified that the subjects had no reported color vision deficiencies and exhibited good color discrimination as assessed with a Farnsworth–Munsell 100 Hue Color Vision Test, with “Superior” or “Average” scores (Total Error Score not exceeding 100). The test was first performed binocularly and then monocularly for the right eye, which was the eye tested in the experiments. For the right eye, we then utilized a minimum motion judgment (Anstis & Cavanagh, 1983; Mullen, et al., 2003; Yoshizawa et al., 2003; Jiang et al., 2022) to determine the isoluminant R-G ratio for each subject at the same visual field location as the stimuli in the main experiments. All subsequent measurements were performed using each subject’s isoluminant R-G ratio. All experiments were performed in a darkened room.

In most of the experiments, subjects reported the envelope motion direction for achromatic or chromatic CM stimuli presented for 250 ms (Figure 2A). Subjects indicated the perceived direction by pressing a key, with subsequent feedback (visual icon on the CRT monitor) for incorrect responses. Envelope direction discrimination performance was measured in a 2AFC task in which subjects reported whether the envelope appeared to move leftwards or rightwards. Based on our previous work with CM stimuli (Ramirez et al., 2022), we fixed the envelope SF at 0.50 cpd, envelope TF at 3 Hz, carrier contrast at 80%, carrier SF at 4 cpd (unless otherwise noted), and carrier TF at 10 Hz. Example stimuli are depicted in Figure 2B for three values of achromatic noise superimposed on either achromatic (top row) or chromatic (bottom row) CM stimuli.

**Figure 2.**
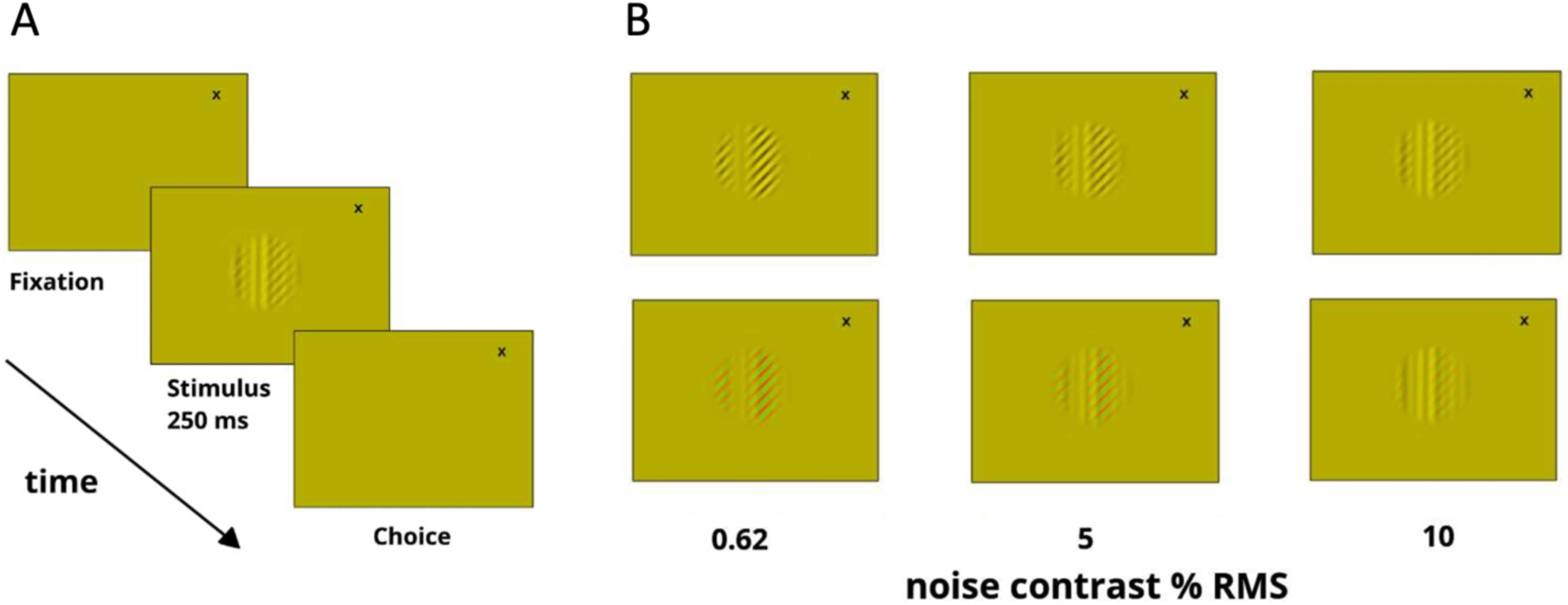
Envelope direction discrimination task and masking noise. A) Direction discrimination task. With fixation on the target (“x”) in the upper right quadrant of the monitor, a CM stimulus appeared for 250 ms. The subject then responded whether the envelope appeared to move leftwards or rightwards (2AFC). The CM stimuli were either achromatic (Y-B) as shown here, or chromatic (R-G). B) Examples of 1-D vertical achromatic (Y-B) masking noise superimposed on CM stimuli. Top row, achromatic (Y-B) CM stimuli, with three levels of noise contrast. Bottom row, same as top row, but for chromatic (R-G) CM stimuli. For purposes of illustration, the SFs shown here are not at the same scale as used in the experiments, and the mixtures of CM stimuli and noise are produced here by graphical transparent overlay.

Using the method of constant stimuli, we tested 5 envelope contrast values logarithmically spaced at 25%, 35%, 50%, 71%, and 100% within each block of trials, with semi-random ordering of trial blocks for achromatic and chromatic CM stimuli. Up to six masking noise contrasts (0.62%, 1.25%, 2.5%, 5%, 6.5%, and 10%) were tested in separate trial blocks arranged in a semi-randomized order. A minimum of three blocks per condition were conducted, resulting in at least 60 trials for each condition.

In addition, detection thresholds were measured for the Y-B noise that was used as the mask. The noise detection task was 2-interval forced-choice (2IFC), with two temporal intervals of 250 ms each separated by a blank period of 250 ms (Figure 3A). The Y-B noise was presented in one of the two intervals at one of a series of noise contrasts, while the other interval was blank. Subjects reported on each trial whether the Y-B noise appeared in the first or second interval.

**Figure 3.**
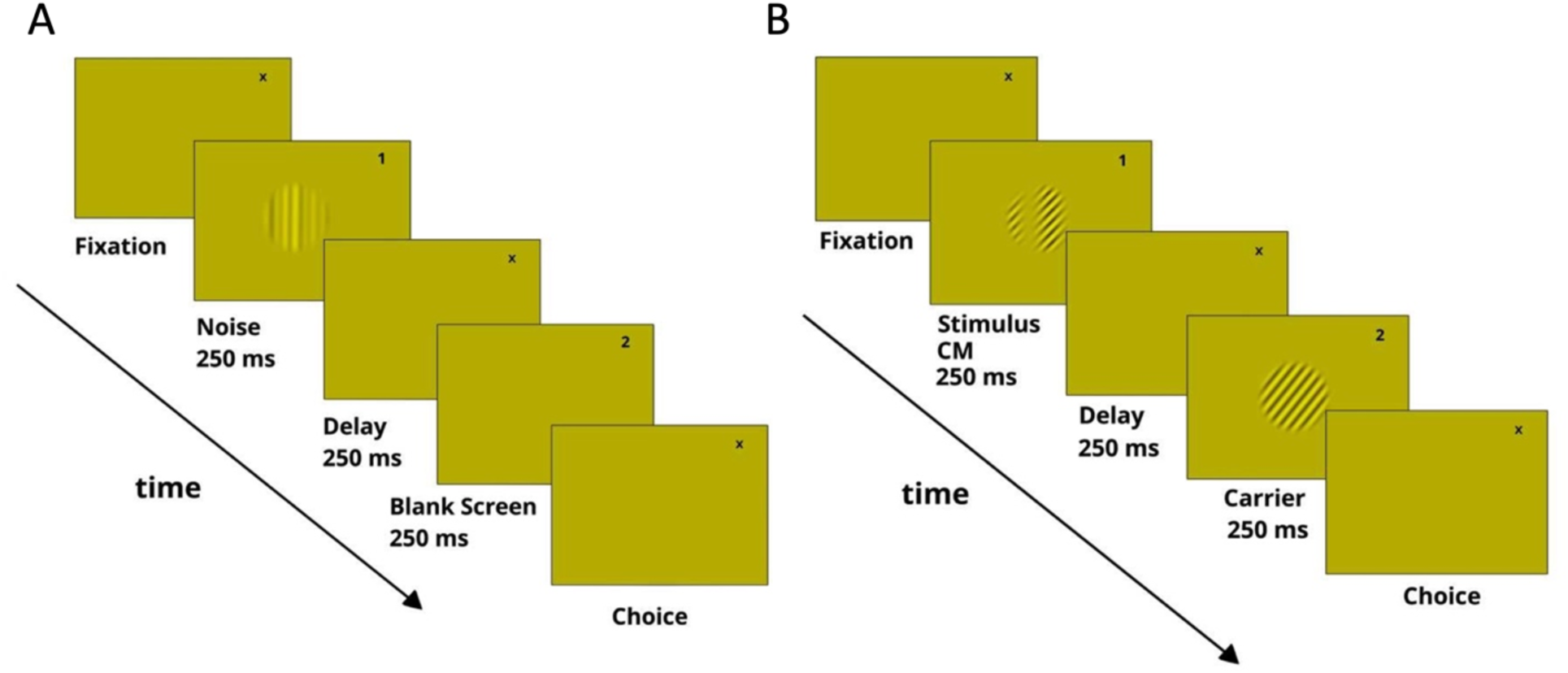
Measurement of detection thresholds. A) Measurement of the detection threshold for the noise mask (achromatic, Y-B) alone, using a 2-interval forced-choice task. A fixation target appeared in the upper right quadrant of the screen: a “1” or “2” during the first or second interval, respectively, and otherwise an “x”. The two intervals (250 ms each) were separated by a blank period (250 ms). A noise stimulus appeared in one of the two intervals (here, the first), while only the fixation target appeared in the other interval. Subjects responded whether the noise appeared in the first or second interval. B) Same as A, but for the measurement of detection thresholds for achromatic (Y-B) and chromatic (R-G) CM stimuli. A CM stimulus was presented in one of the two intervals (here, the first), while an unmodulated carrier was shown in the other. Detection thresholds were measured at varying levels of masking noise contrast (not shown here, for clarity of illustration). The subject responded whether the CM stimulus appeared in the first or second interval.

Detection thresholds were also measured for the achromatic and chromatic CM stimuli at varying levels of masking noise contrast. The detection task was 2IFC, with two intervals of 250 ms each separated by a blank period of 250 ms (Figure 3B). The CM stimulus was presented in one of the two intervals, while the unmodulated carrier alone was shown in the other interval. Subjects reported on each trial whether the CM stimulus appeared in the first or second interval.

### Experimental design and statistical analysis

Except as noted, psychometric functions of percent correct versus a stimulus parameter (envelope contrast for CM stimuli, or RMS contrast for noise) were constructed. For each psychometric function, we determined the best-fitting logistic function and took a threshold value corresponding to the 75% correct level (Figure 4). A bootstrap procedure (Efron & Tibshirani, 1974) was used to estimate the standard error of the threshold. The curve-fitting and bootstrap analysis (400 samples) was performed using the Psychometric Function Fitting of Palamedes Toolbox (Prins & Kingdom, 2018). For cases in which the average performance was below 75% correct for all tested contrasts (e.g., Figure 4, RMS = 5.0), the threshold value was nominally taken as 100% (stars in Figures 6, 7, and 9) for visualization and subsequent curve fitting as described next.

**Figure 4.**
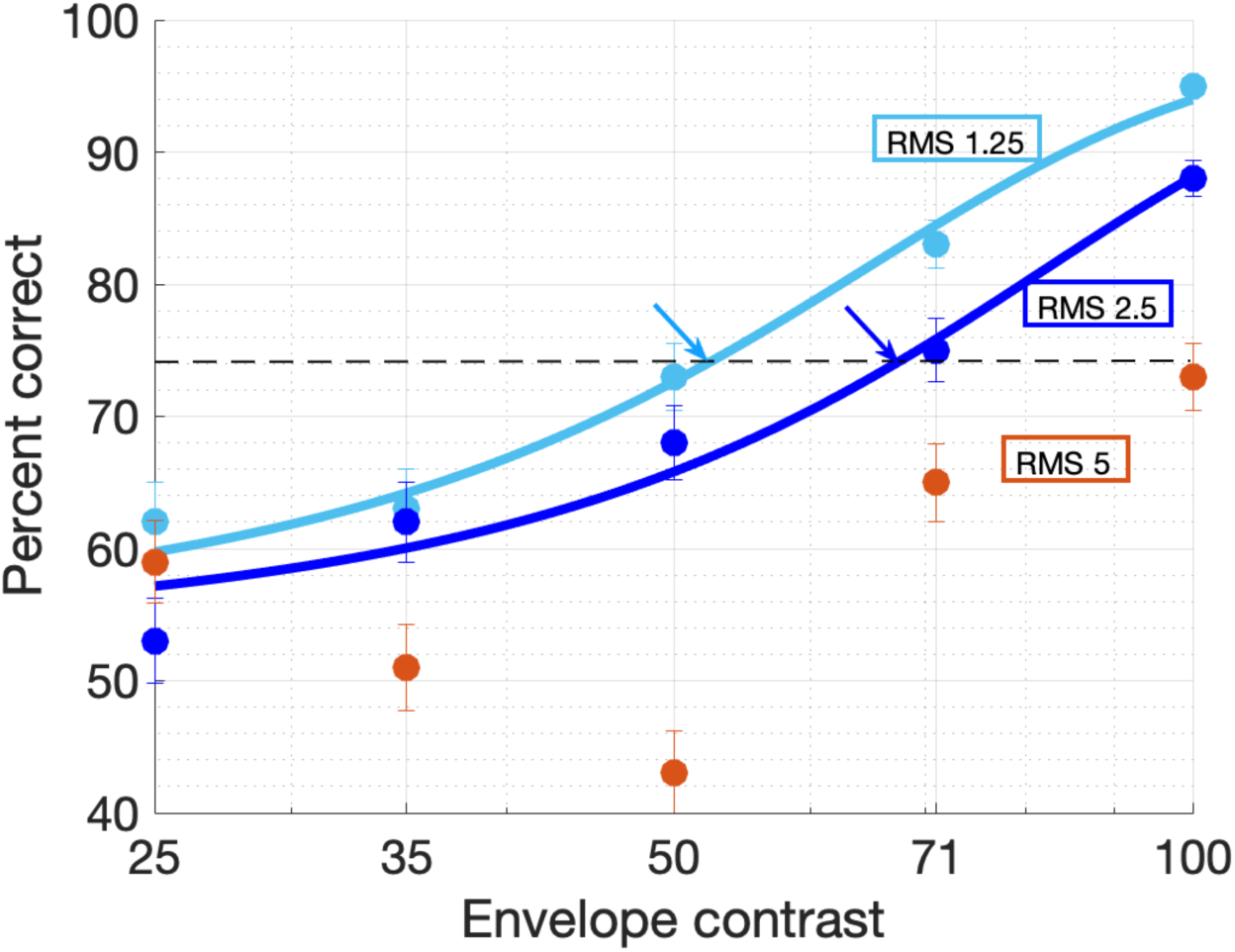
Example psychometric functions and fitted logistic curves. Percent correct versus chromatic (R-G) CM envelope contrast for Subject S3 at 1.25% RMS contrast noise is shown in light blue, at 2.5% RMS contrast in dark blue, and 5% RMS contrast in orange. Error bars indicate the binomial standard error for each condition. The best-fitting logistic function and a threshold value corresponding to the 75% correct level (marked by arrows) were determined for the 1.25% and 2.5% noise contrasts, while for 5% the performance was too poor to provide a curve-fit and threshold.

Plots of envelope contrast threshold versus noise masking contrast (see Results) were typically flat for low noise contrasts up to a point and then systematically increased for larger noise values. To quantify these results, we fit each plot with a descriptive function:

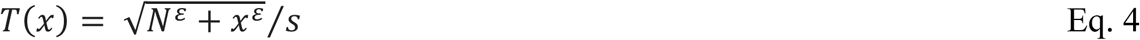

where *T* was the CM envelope contrast threshold, *x* was the masking noise contrast, and *N*, *ε*, and *s* were fitted parameters. The primary parameter of interest was *N*, a “noise effect threshold”, which corresponds to the value of noise contrast at which the CM envelope contrast thresholds began to rise.

## Results

### Experiment 1: Comparison of carrier SF dependence for achromatic and chromatic CM stimuli

Based on previous studies implicating Y-like/parasol retinal ganglion cells in the processing of CM stimuli at relatively high spatiotemporal carrier frequencies (Gharat & Baker, 2017; Rosenberg & Issa, 2011; Ramirez et al., 2022), we initially hypothesized that direction discrimination would be superior for achromatic CM stimuli compared to chromatic CM stimuli. This hypothesis stemmed from the fact that Y-like cells, particularly parasol cells, primarily process achromatic signals, whereas chromatic signals are predominantly processed by midget cells.

A key finding from our previous work with CM stimuli like those used in this study but presented in grayscale (Ramirez et al., 2022) was a bandpass dependence on carrier SF, which provided a signature of a mechanism rooted in Y-like responses. As an initial test with the present stimuli, we compared the dependence on carrier SF for achromatic and chromatic CM stimuli in four subjects using a 2AFC task in which the motion direction of the envelope was reported. In this experiment, we measured percent correct with a fixed carrier contrast of 80%, an envelope contrast of 60%, and a carrier TF of 10 Hz. We varied the carrier SF using a method of constant stimuli across seven levels: 1.4, 2.0, 2.9, 4.2, 5.9, 8.4, and 12.5 cpd.

For most subjects (Figure 5A-C), there was a clear bandpass dependence on carrier SF, although one subject (S4; Figure 5D) exhibited a low-pass dependence. The average across the four subjects’ data was bandpass (Figure 5E). For comparison, the previous results with grayscale CM stimuli are shown in Figure 5F. Similar to the current data, most subjects showed a bandpass carrier SF dependence (Figure 5F). Collectively, these results guided our selection of a carrier SF of 4 cpd for the subsequent experiments.

**Figure 5.**
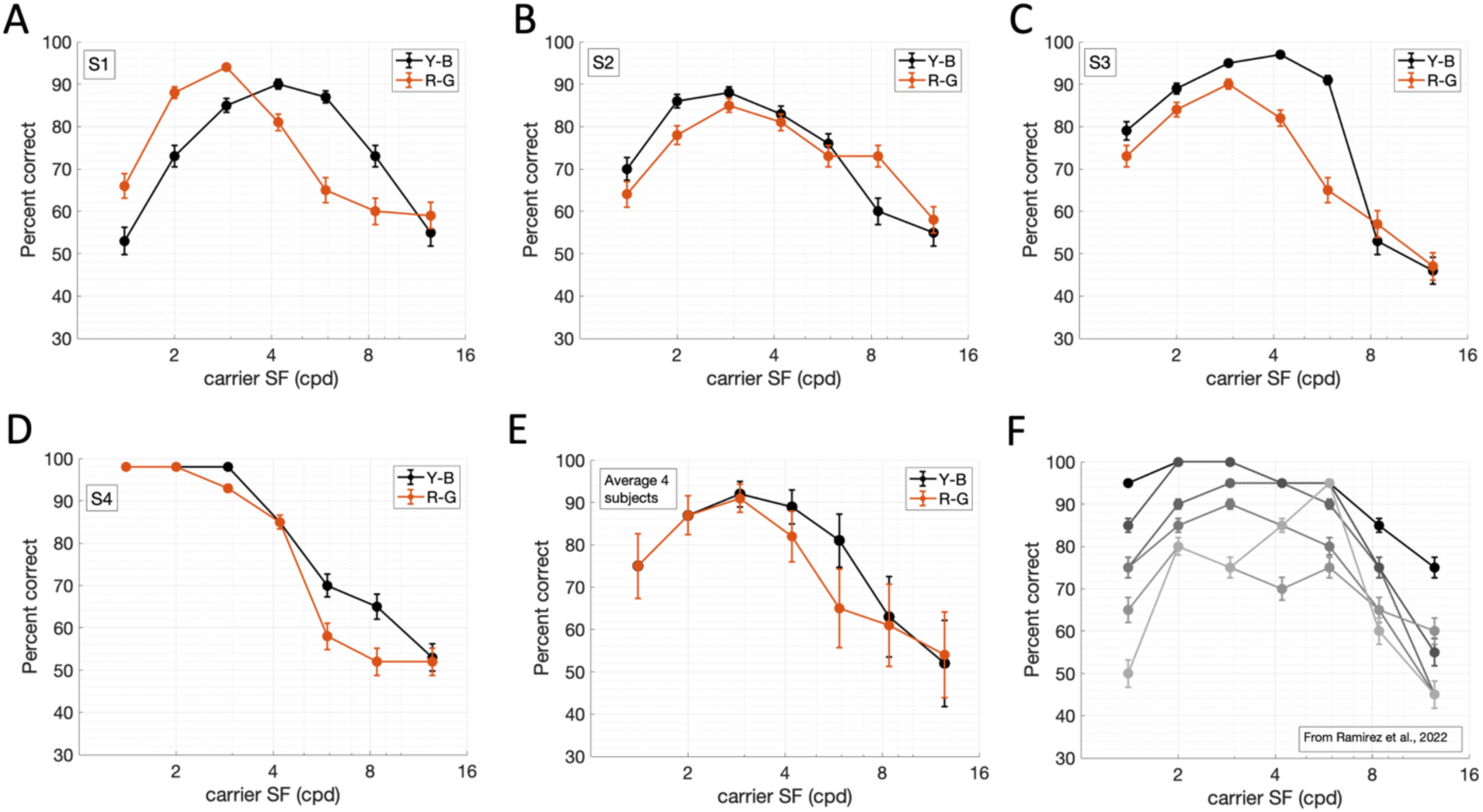
Envelope motion direction discrimination with achromatic (Y-B) and chromatic (R-G) CM stimuli as a function of carrier SF. A-D) Performance (percent correct) with achromatic (Y-B) CM stimuli (black) and chromatic (R-G) CM stimuli (orange) for four subjects (S1-S4). E) Average performance of the four subjects. F) Carrier SF dependence results with grayscale CM stimuli for six subjects from Ramirez et al. (2022). Same stimulus parameters as this study, except that the envelope contrast was 80% rather than 60%. Error bars in all except E indicate binomial standard error (SE) for each condition. For E, error bars indicate SEs across the percent correct values for each of the 4 subjects.

In an apparent conflict with our initial expectations, we found that the chromatic results were generally similar to the achromatic results for most subjects (Figure 5A-E). We therefore wanted to evaluate whether responses to the chromatic CM stimuli may have been mediated by the activation of luminance-based mechanisms, notwithstanding that the stimuli themselves were isoluminant (Mullen et al., 2003).

### Experiment 2: Effect of luminance masking noise on envelope motion direction discrimination for achromatic and chromatic CM stimuli

The similarity in performance between achromatic and chromatic CM stimuli in Experiment 1 may have resulted from the activation of luminance-based mechanisms due to optical chromatic aberrations and/or temporal delays between the processing of long- and middle-wavelength cone signals (Mullen et al., 2003) (see Discussion). If performance with the chromatic stimuli was mediated by activation of a luminance-based mechanism, then addition of spatially and temporally broadband luminance noise (Gegenfurtner & Kiper, 1992; Losada & Mullen, 1995; Yoshizawa et al., 2003) should impair performance much more for chromatic than achromatic motion stimuli. To test this, we introduced spatially 1-D, vertically oriented achromatic masking noise within the cosine-tapered circular window where the stimuli were presented (see Methods).

Performance was assessed in the presence of masking by interleaving on alternate stimulus frames either achromatic or chromatic CM stimuli, and luminance noise (achromatic, Y-B). We tested up to six conditions with different masking noise contrasts (0.62%, 1.25%, 2.5%, 5%, 6.5%, and 10%), measured as RMS contrast (see Methods), in separate trial blocks in a semi-randomized order. The envelope SF was 0.50 cpd, envelope TF was 3 Hz, carrier contrast was 80%, carrier SF was 4 cpd, and carrier TF was 10 Hz.

We first examined performance with achromatic CM stimuli for six subjects. In each panel of Figure 6, the envelope contrast threshold for envelope motion discrimination is plotted as a function of masking noise contrast. For each subject, performance was fairly robust to the level of masking noise contrast until around 5-6%. At higher levels, performance systematically deteriorated, culminating in an inability to determine a threshold (indicated by a star symbol, placed nominally at 100% – see example at 5% RMS contrast in Figure 4).

**Figure 6.**
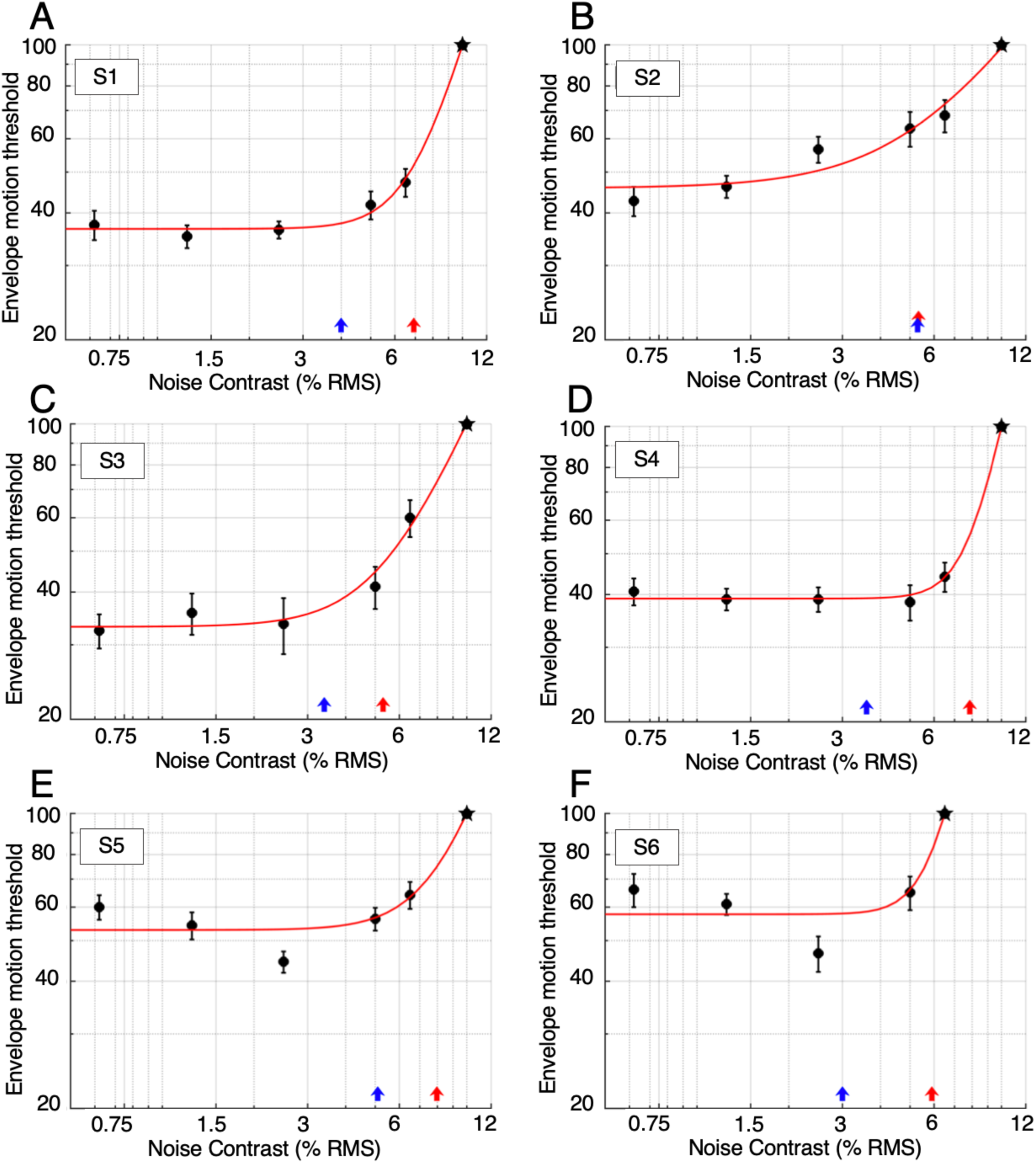
Thresholds for envelope motion direction discrimination with achromatic (Y-B) CM stimuli masked by luminance (achromatic, Y-B) noise. Each plot shows, for one subject, the envelope contrast motion direction discrimination threshold as a function of masking noise contrast. Star symbols indicate performance too poor to fit a psychometric function, here plotted nominally at 100% (e.g., see data for 5% RMS noise contrast in Figure 4). Error bars indicate standard errors of the thresholds estimated with a bootstrap procedure. Red curves are fitted functions (Equation 4) used to estimate noise effect thresholds indicating the level of noise contrast necessary to impair performance (red arrows). The blue arrows along the abscissa mark the subjects’ noise detection thresholds.

Notably, the marked decline in discrimination performance with increasing noise levels did not begin until the noise level was above its detection threshold, indicated by a blue arrow on each plot (Figure 6), with the possible exception of Subject S2. This indicates that, for achromatic CM stimuli, the noise began to exert a masking effect only after it had become visible.

Results are shown for the same subjects with chromatic CM stimuli in Figure 7. While the chromatic CM results followed a similar overall trend as their achromatic counterparts, the measured thresholds were often higher for the chromatic CM stimuli (Figure 7) than for the achromatic CM stimuli (Figure 6). To quantify this difference, we performed a pairwise comparison of the 29 cases (subjects, noise contrasts) for which a measurable threshold could be obtained for at least one of the compared values. Non-measurable thresholds (as in e.g., Figure 4, RMS = 5.0), were nominally taken as 100%. A Wilcoxon signed-rank test showed that achromatic envelope motion thresholds (Mdn = 44.5) were significantly lower than corresponding thresholds for chromatic motion (Mdn = 71), W = 1.00, z = –4.60, *p* < 0.001.

**Figure 7.**
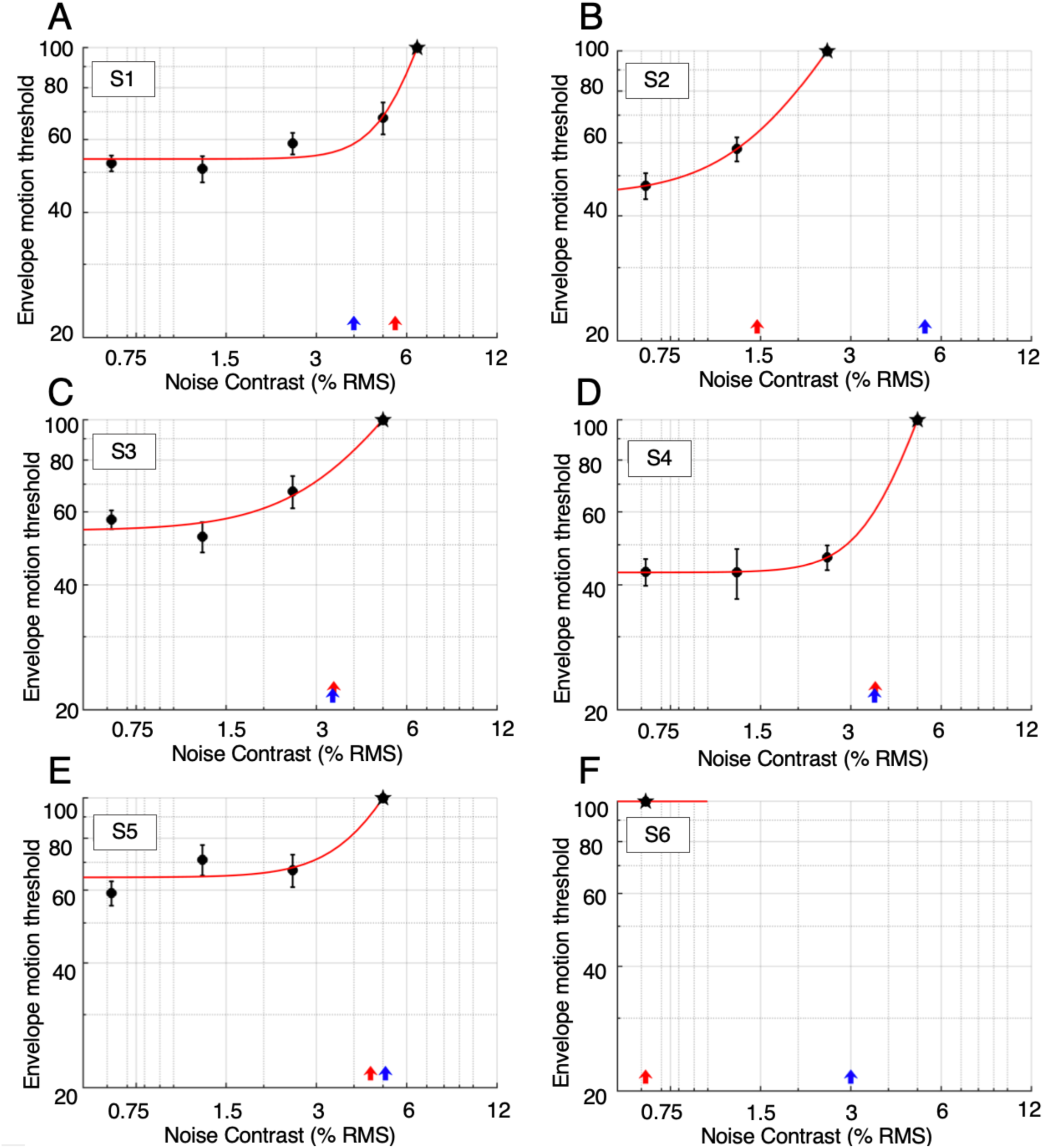
Thresholds for envelope motion direction discrimination with chromatic (Y-B) CM stimuli masked by luminance (achromatic, Y-B) noise, shown as in Figure 6. Subject S6 was unable to perform the task even at the lowest noise contrast tested.

Moreover, noise contrast affected performance with the chromatic CM stimuli to varying degrees across the subjects. At one end of that spectrum, Subject S6 (Figure 7, panel F) was unable to perform the task sufficiently to provide psychometric functions, even at the lowest masking noise contrast of 0.62%. Subject S2 (panel B) exhibited good performance at noise contrasts up to 1.25%, while Subjects S3 (panel C), S4 (panel D), and S5 (panel E), performed well up to 2.5%. On the other end of that spectrum, Subject S1 (panel A) maintained relatively good performance at masking noise contrasts up to 5%.

For the chromatic CM stimuli, performance typically began to decline at noise contrast levels near or below the point at which the noise became detectable (blue arrows in Figure 7). This pattern differed markedly from that with achromatic CM stimuli, for which performance did not decay until after the noise was detectable. The negative impact of undetectable masking noise on chromatic stimuli is consistent with the spurious activation of a luminance-based mechanism due to optical chromatic aberrations or differential temporal dynamics for long- and middle-wavelength cone signals (Gegenfurtner & Kiper, 1992; Losada & Mullen, 1995; Mullen et al., 2003; Yoshizawa et al., 2003).

To quantify the relationship between the effect of masking noise on motion discrimination thresholds and the noise detection thresholds, we first obtained quantitative estimates of the knee points of the above plots, using curve fits to the data in Figures 6 and 7. The functions (Equation 4) were flat at low noise contrasts and began to increase at a knee point corresponding to the parameter *N*, which we took as an estimate of the “noise effect threshold” (red arrows in Figures 6 and 7). This parameter provides a way to compare the relationship between the minimum masking noise necessary to affect the motion discrimination performance and the noise detection threshold (blue arrows in Figures 6 and 7).

The results of this comparison for achromatic stimuli are shown in Figure 8A. The noise effect thresholds (M = 6.7, SD = 1.1) were greater than the noise detection thresholds (M = 4.0, SD = 0.4), and a paired t-test confirmed that the difference was significant: t(5) = 4.5, *p* = 0.006. This finding indicates that the noise exerted a masking effect only at contrasts at which it was readily visible. For chromatic stimuli (Figure 8B), the noise effect (M = 3.1, SD = 1.8) appeared at a contrast near or below the detection threshold for most subjects (M = 4.0, SD = 0.9), and a t-test indicated that the difference was not significant: t(5) = -1.2, *p* = 0.28. For Subjects S2 and S6, noise masking for chromatic stimuli occurred at contrasts far below the detection threshold. This quantification confirms that chromatic motion was masked by luminance noise that was, at most, barely detectable, and in some cases not detectable at all to the subject.

**Figure 8.**
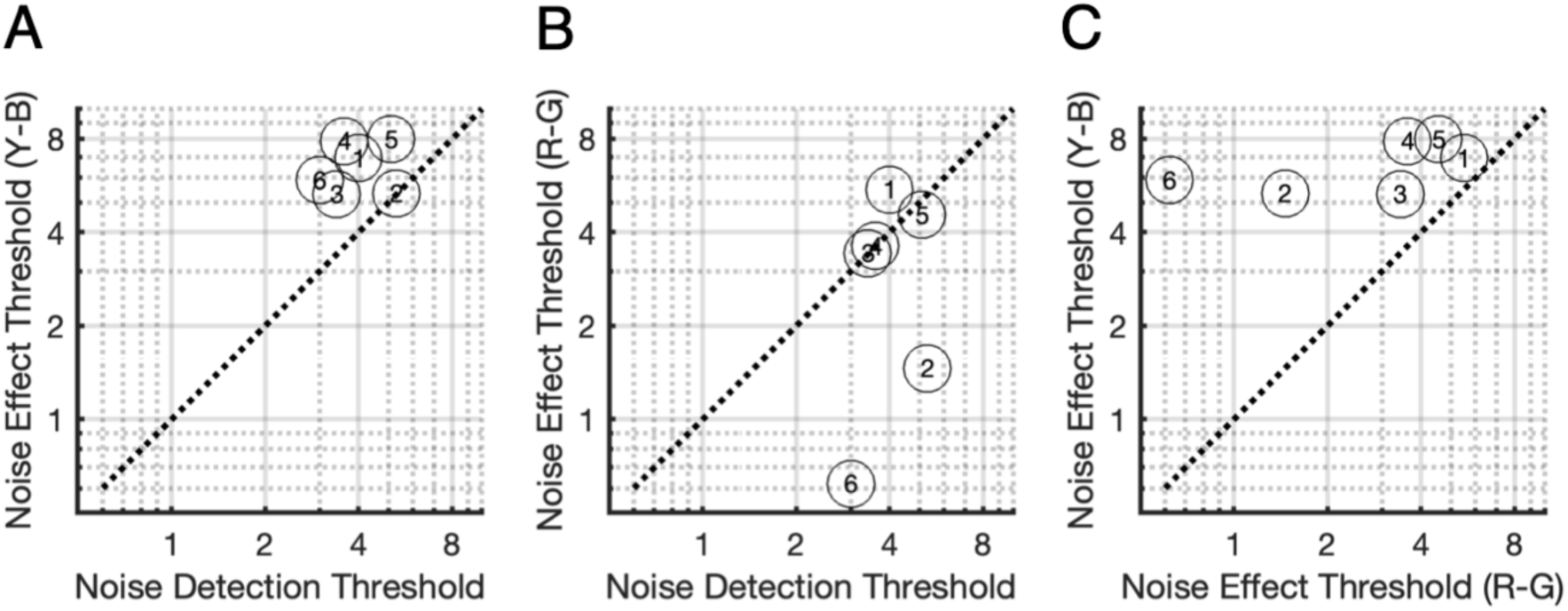
Efficacy of masking noise in relation to its detection threshold. A) Noise effect thresholds for each of six subjects, estimated from curve-fits (Figure 6), as a function of the noise detection thresholds for achromatic (Y-B) envelope motion discrimination. B) Same as A but for the chromatic (R-G) CM stimuli (Figure 7). C) Relation between noise effect thresholds for achromatic versus chromatic CM stimuli. Diagonal dotted line indicates 1:1 relationship.

To further evaluate these differences, we compared the achromatic and chromatic noise effect thresholds. The noise effect thresholds for the achromatic CM stimuli (M = 6.7, SD = 1.09) were higher than those for the chromatic CM stimuli (M = 3.1, SD = 1.8) across all subjects (Figure 8C), and the difference (M = 3.5, SE = 0.57) was significant with a paired t-test: t(5) = 6.2, *p* < 0.001. These results collectively indicate that chromatic CM motion was substantially more vulnerable to masking by luminance noise than achromatic CM motion, consistent with the idea that chromatic CM motion is mediated by a luminance-based mechanism.

### Experiment 3: Envelope detection thresholds for chromatic CM stimuli

It is conceivable that the larger effect of noise masking on chromatic than achromatic CM stimuli might have arisen due to a poor ability to reliably see the chromatic CM envelope. To address this possibility, we measured detection thresholds for the envelopes of chromatic CM stimuli with three of the subjects, again at varying levels of achromatic masking noise (Figure 9). The CM detection thresholds were relatively stable at lower noise contrasts, before increasing. Importantly, in each case, the envelope detection performance (red arrows) exceeded the noise detection threshold performance (blue arrows). Specifically, Subject S1 was able to perform the envelope detection task at noise levels up to 6.5% (Figure 9A) whereas envelope motion discrimination declined beyond 5% (Figure 7A), Subject S3 was able to perform the detection task up to 10% noise (Figure 9B) whereas discrimination declined beyond 2.5% (Figure 7C), and Subject S4 was able to detect the envelope up to 6.5% noise (Figure 9C) whereas discrimination declined beyond 2.5% (Figure 7D). These results indicate that poorer performance with chromatic CM stimuli was not due to an inability to detect the envelope, but rather an impairment in discriminating its motion. Because the luminance noise effectively masked envelope direction discrimination for chromatic CM stimuli but exerted a relatively small impairment in detection, these results further point to the contribution of luminance-based mechanisms in envelope motion perception.

**Figure 9.**
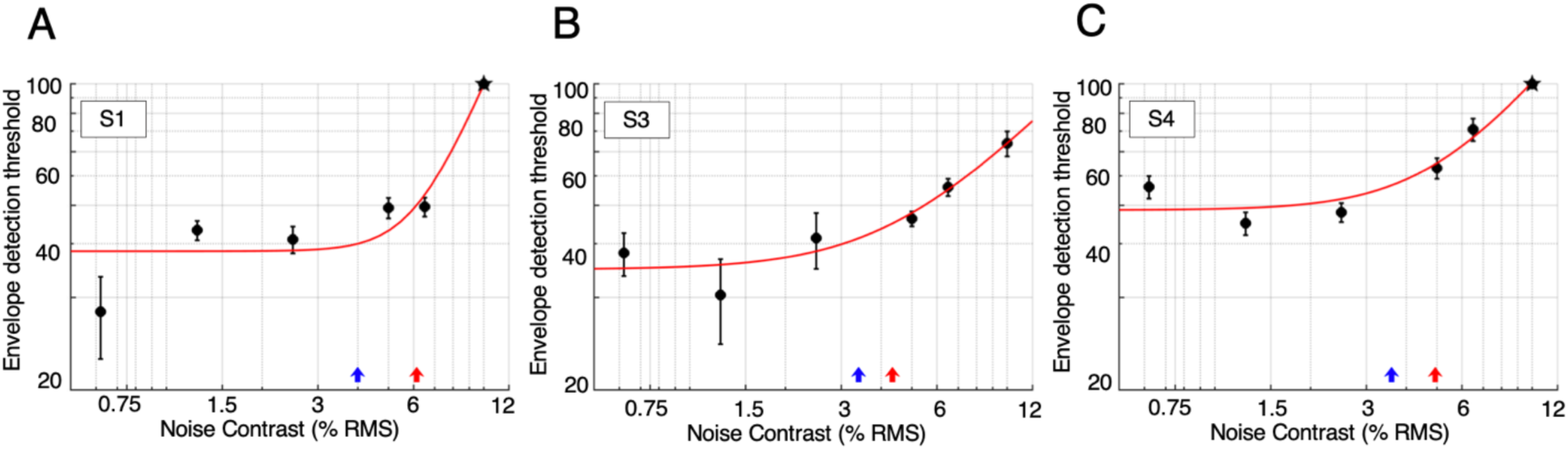
Envelope detection thresholds for chromatic (R-G) CM stimuli. Each plot shows detection thresholds for CM stimuli as a function of RMS noise contrast, for varying levels of achromatic masking noise. Error bars indicate estimated standard errors of the thresholds. Red curves are fitted functions (Equation 4) used to estimate noise effect thresholds indicating the level of noise contrast necessary to impair performance. Red arrows mark the corresponding noise effect thresholds. Blue arrows mark the subjects’ noise detection thresholds.

Lastly, for a direct quantitative comparison of detection and direction discrimination performance for the envelope of chromatic CM stimuli, we employed curve-fits as previously to measure noise effect thresholds for detection. Figure 10A shows that for all three subjects, the noise masking effect for detection occurred at contrasts above the noise detection thresholds – that is, the noise exerted a masking effect only when it was visible. Likewise, Figure 10B shows that the noise masking effect for detection thresholds was always higher than for direction discrimination – thus chromatic direction discrimination was more sensitive than detection to luminance noise masking. These findings reinforce the conclusion that luminance-based mechanisms support the perception of chromatic envelope motion.

**Figure 10.**
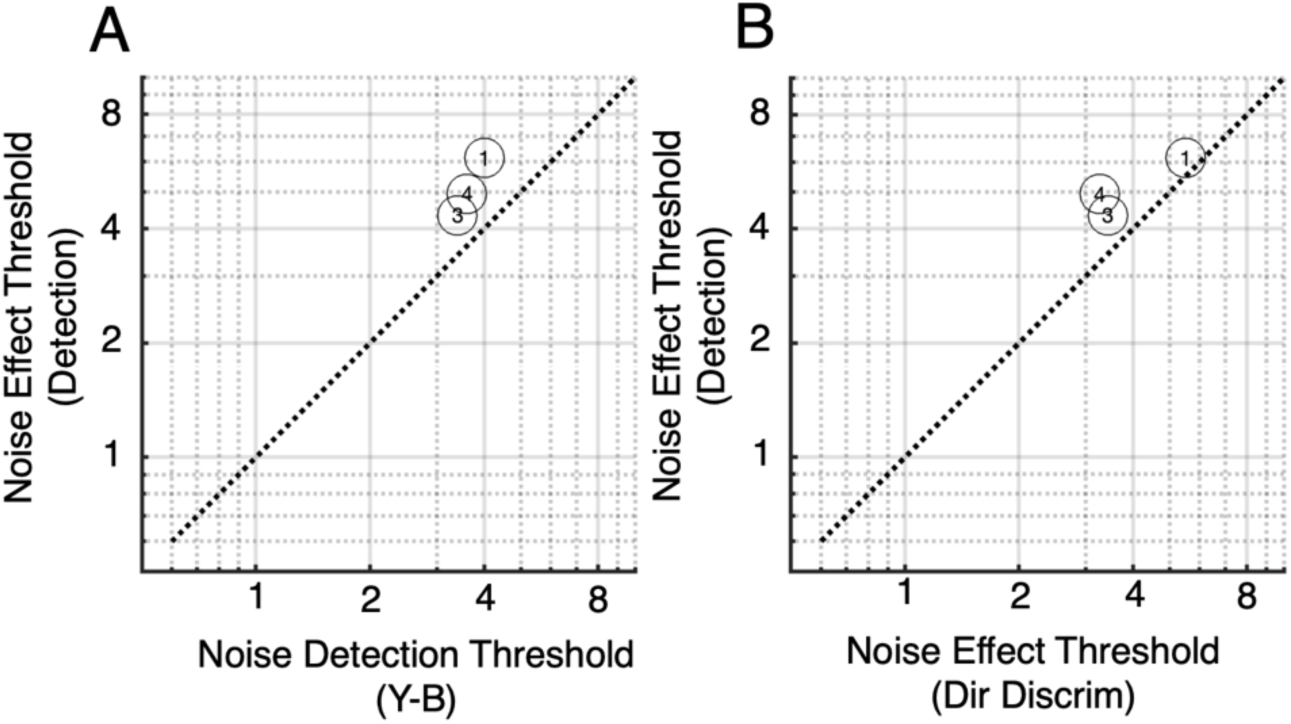
Efficacy of masking noise on detection thresholds for chromatic (R-G) CM stimuli. A) Noise effect thresholds for detection of chromatic (R-G) CM stimuli for three subjects, estimated from curve-fits (Figure 9), as a function of the noise detection thresholds. B) Same as A but as a function of noise effect thresholds for direction discrimination with chromatic (R-G) CM stimuli. Diagonal dotted line indicates 1:1 relationship.

## Discussion

In this study, we used CM stimuli designed to activate the nonlinear properties of Y-like/parasol cells in the magnocellular pathway by presenting stimuli at high spatiotemporal carrier frequencies (Zhou and Baker, 1994; Rosenberg & Issa, 2011; Ramirez et al., 2022). To assess whether the CM stimuli selectively activated Y-like cells, we evaluated motion direction discrimination performance for both achromatic (Y-B) and chromatic (R-G) CM stimuli. Additionally, we introduced luminance masking noise to disrupt luminance-based mechanisms of motion processing arising from chromatic aberrations. Chromatic CM motion was notably more vulnerable to masking by luminance noise than achromatic CM motion, suggesting that envelope motion perception relied on luminance signals. Critically, this vulnerability did not stem from difficulties in detecting the CM envelope itself but rather from challenges in discerning its motion. Indeed, luminance noise effectively disrupted envelope direction discrimination for chromatic CMs yet had only a relatively minor impact on its detection. These results imply that envelope motion perception with chromatic CM stimuli is enabled by luminance mechanisms.

### Dependence on carrier spatial frequency

Drawing on neurophysiology research implicating Y-like/parasol retinal ganglion cells in cortical responses to CM stimuli at high spatiotemporal carrier frequencies (Gharat & Baker, 2017; Rosenberg & Issa, 2011), we predicted that psychophysical direction discrimination would be better for achromatic than chromatic CM stimuli. This prediction was based on the fact that

Y-like/parasol cells primarily process achromatic signals, while chromatic (R-G) signals are mainly processed by midget cells (Shapley & Perry, 1986; Nassi & Callaway, 2009). In Experiment 1, our findings with CM stimuli (Figure 5) mirrored previous results from Ramirez et al. (2022) with grayscale CM stimuli, demonstrating a bandpass dependence on carrier SF. This bandpass characteristic for relatively high carrier SFs supported the mediation by a CM mechanism associated with Y-like responses.

The similarity of our findings with achromatic and chromatic CM stimuli (Figure 5) may appear contradictory to the idea that CM performance is mediated by Y-like cells in the magnocellular pathway, which responds best to achromatic stimuli. However, our subjects reported that the chromatic CM stimuli seemed less visible and/or that their motion was less clear than their achromatic counterparts. These subjective reports and previous findings on luminance contamination in motion of chromatic stimuli (Mullen et al., 2003; Yoshizawa et al., 2003) motivated the use of luminance masking noise in our experiments.

### Effects of luminance masking noise on CM motion discrimination

Studies investigating the perception of motion with chromatic isoluminant stimuli have yielded varied findings. In some cases, the motion was perceived as slower, or to momentarily disappear, compared to luminance-based stimuli (Cavanaugh et al., 1984; Livingstone & Hubel, 1988; Mullen & Boulton 1992a,1992b). Another study observed a complete loss of apparent motion perception with random dot kinematograms (Ramachadran & Gregory, 1978). However, other evidence indicates that motion tasks can be performed under isoluminant conditions (Cropper & Derrington, 1996; Dobkins & Albright, 1993). This ability may be mediated by mechanisms other than chromatic processing, potentially involving responses from the luminance pathway rather than a specialized chromatic mechanism. For instance, luminance-based mechanisms could be activated by chromatic aberrations arising from optical imperfections (Bradley et al., 1992; Flitcroft, 1989) or delays between the responses of long- and middle-wavelength cones as they are integrated into a luminance pathway (Mullen et al., 2003; Yoshizawa et al., 2000). This likely occurs early in the visual system (Stromeyer et al., 1995, 1997, 2000; Tsujimura et al., 1999, 2000). In such cases, demonstrating that a superimposed luminance noise mask selectively impairs chromatic motion performance can imply the contamination of chromatic motion processing by luminance signals (Mullen et al., 2003; Yoshizawa et al., 2000, 2003).

In Experiment 2, we followed this logic by introducing spatially and temporally broadband luminance noise to interfere with (mask) spurious luminance signals. Our results revealed a significant impairment in motion discrimination for chromatic stimuli compared to achromatic stimuli. As expected, for both types of CM stimuli, discrimination performance declined as noise contrast increased, ultimately resulting in an inability to determine a threshold. Critically, for achromatic CM stimuli, the decline in performance began after noise levels exceeded the noise detection threshold, indicating that the masking effect occurred only when the noise was visible. On the other hand, for chromatic CM stimuli, the decline in performance began at noise levels which were at most barely detectable, and in some cases not at all detectable. Indeed, the noise effect thresholds for achromatic CM stimuli were greater than those for chromatic CM stimuli across all subjects (Figure 8C). Taken together, these results suggest that chromatic CM motion was considerably more susceptible to masking from luminance noise than achromatic CM motion, consistent with the idea that chromatic CM motion is processed via a luminance-based mechanism.

The impact of masking noise on chromatic stimuli also varied considerably among subjects, in comparison to its relatively uniform impact on achromatic stimuli. Intersubject differences with chromatic stimuli might be explained by variations in optical imperfections including longitudinal chromatic aberration (LCA) and transverse chromatic aberration (TCA), both of which result from dispersion within the eye’s refractive components. LCA occurs along the optical axis of the lens, where different wavelengths of light focus at different distances, leading to variations in refractive power with wavelength. Shorter wavelengths exhibit a higher refractive index and greater refractive power, resulting in LCA of approximately two diopters across the visible spectrum (Howarth & Bradley, 1986; Rynders, Navarro, and Losada, 1998). TCA reflects differences in the angular position of the retinal image for various wavelengths, being particularly pronounced in decentered pupils (Thibos, Bradley & Zhang, 1991; Walsk, 1988), and increasing with retinal eccentricity (Ogboso & Bedell, 1987; Thibos, 1987). TCA can further differ across subjects due to variations in foveal position relative to the optical axis (Le Grand, 1956) and pupil size (Marcos et al., 1999). Cylindrical defects such as astigmatism amplify the effects of TCA, which is more directly influenced by the overall refractive structure of the eye. While astigmatism can also indirectly worsen defocus, its impact on LCA is less pronounced (Thibos, 1987). As discussed earlier, relative delays in the L- and M-cone signals can also generate substantial luminance mechanism signals from chromatic stimuli. Variations across subjects in such “temporal chromatic aberrations” (Mullen et al., 2003) are not well documented, but could also underlie some of the intersubject differences we found in noise masking effects. Thus, the extent to which chromatic aberrations introduce spurious luminance signals is expected to vary across subjects, and this may have contributed to intersubject variability in the masking effects we observed with chromatic CM stimuli. Nevertheless, our finding that chromatic CM motion was more susceptible to masking by luminance noise than achromatic CM motion supports the idea that its perception is mediated by a luminance mechanism.

### Is impaired motion simply due to impaired detection?

In Experiment 3, we investigated whether poorer motion performance with chromatic than achromatic CM stimuli might simply arise from a diminished ability to detect the stimuli. To address this question, we measured detection thresholds for the envelopes of chromatic CM stimuli at different levels of masking noise. Importantly, the procedure to measure CM detection thresholds ensured that the performance reflected detection of the CM’s envelope, and not simply detection of the carrier. This was achieved by employing a two-interval choice task in which the carrier was presented in both intervals, with contrast-modulation by an envelope only in the target interval. The results verified that poorer performance with chromatic CM stimuli was not due to an inability to detect the envelope, but rather to difficulties in discriminating its motion. These findings suggest that luminance mechanisms play a predominant role in processing the motion of chromatic (R-G) CM stimuli, as luminance noise effectively masked their envelope direction discrimination while causing minimal impairment in detection (Mullen et al., 2003).

### Neural mechanisms

Early research in the cat retino-geniculate pathway identified two main types of RGCs: X cells and Y cells (Enroth-Cugell & Robson, 1966). Although Y cells were initially discovered in cats, they have counterparts in a wide range of mammals (Peichl, 1991). In primates, the two most numerous RGC types – midget and parasol cells – may correspond to X cells (compact neurons with small receptive fields) and Y cells (neurons with large receptive and dendritic fields), respectively. In particular, primate parasol RGCs give first harmonic (linear) responses to gratings with low spatial frequencies, and second harmonic (nonlinear) responses at high spatial frequencies (Crook et al., 2008), which is characteristic of cat Y-cells (Hochstein & Shapley, 1976). Midget cells are generally associated with the parvocellular pathway, while parasol cells contribute to the magnocellular pathway (Nassi & Callaway, 2009). Despite this overall consensus, there has been some debate. It has been suggested that in the primate LGN, X-like cells are found not only in the parvocellular layers but also in the magnocellular layers, referred to as “Magno-X cells” – whereas Y-like cells were termed “Magno-Y cells” (Kaplan & Shapley, 1982). The primary distinction between Magno-X and Magno-Y cells lay in the nonlinear behavior of Magno-Y cells, like that of cat Y-cells as described in the Introduction. Regardless, our results support the idea that Y-like cells mediate processing of CM stimuli.

In some cases, RGCs from the magnocellular pathway respond, albeit weakly, to red-green modulated gratings. These responses are likely due to a rectified response to a chromatic signal originating from M- and L-cone inputs (Lee & Sun, 2009). Similar effects possibly originating from the magnocellular pathway (Stromeyer et al., 1997), may facilitate luminance-based direction discrimination of moving isoluminant chromatic gratings. Although such mechanisms could have played a role in the observed motion of chromatic CMs at isoluminance, our use of luminance noise masking likely minimized their influence.

### Conclusions

Overall, these experiments support the notion that achromatic CMs could be employed to selectively assess the function of Y-like cells, such as parasol cells, in the magnocellular pathway. This selectivity might be of value for clinical applications, as the magnocellular pathway has been implicated in glaucoma (Quigley et al., 1987, 1988; Glovinsky et al., 1991; Yücel et al., 2003; Shou et al., 2003; Zhang et al., 2016) and dyslexia (Livingstone et al., 1991; Laprevotte 2021). In particular, the development of specific tests of magnocellular function might facilitate the early diagnosis and tracking of such conditions.

## Acknowledgments

Supported by Canadian NSERC (National Science and Engineering Research Council) Discovery grant RGPIN-2023-03559 to CLB and National Institutes of Health Grants (EY029438 & EY035005) to AR.

We thank Ethan Pirso for contributions to stimulus programming. We are also grateful to Kathy Mullen, Fred Kingdom, and Alexander Baldwin for useful comments and advice.

## Notes

**funding:** Canadian NSERC (Natural Science and Engineering Research Council of Canada) Discovery grant RGPIN-2023-03559 to CLB and National Institutes of Health Grants (EY029438 & EY035005) to AR.

### Competing Interest Statement

Based in part on work associated with these findings, the McGill Royal Institution and the Wisconsin Alumni Research Foundation hold patent US20210312613A1 for detecting early-stage glaucoma and optic nerve diseases, with inventors Ari Rosenberg, Curtis Baker, and Ana Ramirez.

## References

Anstis, S., & Cavanagh, P. (1983). A minimum motion technique for judging equiluminance. Academic Press.

Baker Jr, C. L., Boulton, J. C., & Mullen, K. T. (1998). A nonlinear chromatic motion mechanism. Vision Research, 38(2), 291–302.

Bradley, A., Zhang, X., & Thibos, L. (1992). Failures of isoluminance caused by ocular chromatic aberrations. Applied Optics, 31(19), 3657–3667.

Brainard, D. H. (1997). The Psychophysics Toolbox. Spatial Vision, 10, 433–436.

Cavanagh, P., & Mather, G. (1989). Motion: the long and short of it. Spatial vision, 4(2-3), 103–129.

Crook, J. D., Peterson, B. B., Packer, O. S., Robinson, F. R., Gamlin, P. D., Troy, J. B., & Dacey, D. M. (2008). The smooth monostratified ganglion cell: evidence for spatial diversity in the Y-cell pathway to the LGN and superior colliculus in the macaque monkey. The Journal of neuroscience: the official journal of the Society for Neuroscience, 28(48), 12654.

Cropper, S. J., & Derrington, A. M. (1996). Detection and motion detection in chromatic and luminance beats. Journal of the Optical Society of America A, 13(3), 401–407.

Dacey, D. M., & Lee, B. B. (1994). The’blue-on’opponent pathway in primate retina originates from a distinct bistratified ganglion cell type. Nature, 367(6465), 731–735.

Demb, J. B., Haarsma, L., Freed, M. A., & Sterling, P. (1999). Functional circuitry of the retinal ganglion cell’s nonlinear receptive field. Journal of Neuroscience, 19(22), 9756–9767.

Dobkins, K. R., & Albright, T. D. (1993). Color, luminance, and the detection of visual motion. Current Directions in Psychological Science, 2(6), 189–193.

Efron, B., & Tibshirani, R. J. (1994). An introduction to the bootstrap. Chapman and Hall/CRC.

Enroth-Cugell, C. & Robson, J. G. (1966). The contrast sensitivity of retinal ganglion cells of the cat. J Physiol 187, 517–552.

Faubert, J., Bilodeau, L., & Simonet, P. (2000). Transverse chromatic aberration and colour-defined motion. Ophthalmic and Physiological Optics, 20(4), 274–280.

Flitcroft, D. (1989). The interactions between chromatic aberration, defocus and stimulus chromaticity: implications for visual physiology and colorimetry. Vision Research, 29(3), 349–360.

Gegenfurtner, K. R., & Kiper, D. C. (1992). Contrast detection in luminance and chromatic noise. Journal of the Optical Society of America A, 9(11), 1880–1888.

Gharat, A., & Baker Jr, C. L. (2012). Motion-defined contour processing in the early visual cortex. Journal of neurophysiology, 108(5), 1228–1243.

Gharat, A., & Baker Jr, C. L. (2017). Nonlinear Y-like receptive fields in the early visual cortex: An intermediate stage for building cue-invariant receptive fields from subcortical Y cells. Journal of Neuroscience, 37(4), 998–1013.

Glovinsky, Y., Quigley, H., & Dunkelberger, G. (1991). Retinal ganglion cell loss is size dependent in experimental glaucoma. Investigative ophthalmology & visual science, 32(3), 484–491.

Hochstein, S., & Shapley, R. (1976). Linear and nonlinear spatial subunits in Y cat retinal ganglion cells. The Journal of physiology, 262(2), 265–284.

Howarth, P. A., & Bradley, A. (1986). The longitudinal chromatic aberration of the human eye, and its correction. Vision Research, 26(2), 361–366.

Jiang, Z., Shooner, C., & Mullen, K. T. (2022). Achromatic and chromatic perceived contrast are reduced in the visual periphery. Journal of Vision, 22(12), 3–3.

Kaplan, E., & Shapley, R. (1982). X and Y cells in the lateral geniculate nucleus of macaque monkeys. The Journal of physiology, 330(1), 125–143.

Kleiner, M., Brainard, D., Pelli, D., Ingling, A., Murray, R., Broussard, C. (2007). What’s new in Psychtoolbox-3? Perception 36, 1–16.

Laprevotte, J., Papaxanthis, C., Saltarelli, S., Quercia, P., & Gaveau, J. (2021). Movement detection thresholds reveal proprioceptive impairments in developmental dyslexia. Scientific reports, 11(1), 299.

Le Grand, Y. (1956). Optique physiologique: L’espace visue (Vol. III). Éditions de la Revue d’Optique

Lee, B. B., & Sun, H. (2009). The chromatic input to cells of the magnocellular pathway of primates. Journal of Vision, 9(2), 15–15.

Leventhal, A. G., Rodieck, R., & Dreher, B. (1981). Retinal ganglion cell classes in the Old World monkey: morphology and central projections. Science, 213(4512), 1139–1142.

Li, G., Yao, Z., Wang, Z., Yuan, N., Talebi, V., Tan, J., Wang, Y., Zhou, Y., & Baker Jr, C. L. (2014). Form-cue invariant second-order neuronal responses to contrast modulation in primate area V2. Journal of Neuroscience, 34(36), 12081–12092.

Livingstone, M., & Hubel, D. (1988). Segregation of form, color, movement, and depth: anatomy, physiology, and perception. Science, 240(4853), 740–749.

Livingstone, M. S., Rosen, G. D., Drislane, F. W., & Galaburda, A. M. (1991). Physiological and anatomical evidence for a magnocellular defect in developmental dyslexia. Proceedings of the National Academy of Sciences, 88(18), 7943–7947.

Losada, M. A., & Mullen, K. T. (1995). Color and luminance spatial tuning estimated by noise masking in the absence of off-frequency looking. Journal of the Optical Society of America A, 12(2), 250–260.

Marcos, S., Burns, S. A., Moreno-Barriusop, E., & Navarro, R. (1999). A new approach to the study of ocular chromatic aberrations. Vision Research, 39(26), 4309–4323.

Mareschal, I., & Baker Jr, C. L. (1998). Temporal and spatial response to second-order stimuli in cat area 18. Journal of neurophysiology, 80(6), 2811–2823.

Merigan, W. H. (1991). P and M pathway specialization in the macaque. In From pigments to perception: Advances in understanding visual processes (pp. 117–125). Springer.

Merigan, W. H., Byrne, C. E., & Maunsell, J. (1991). Does primate motion perception depend on the magnocellular pathway? Journal of Neuroscience, 11(11), 3422–3429.

Metha, A. B., & Mullen, K. T. (1997). Red–green and achromatic temporal filters: a ratio model predicts contrast-dependent speed perception. Journal of the Optical Society of America A, 14(5), 984–996.

Mullen, K. T., & Baker Jr, C. L. (1985). A motion aftereffect from an isoluminant stimulus. Vision Research, 25(5), 685–688.

Mullen, K. T., & Boulton, J. C. (1992a). Absence of smooth motion perception in color vision. Vision Research, 32(3), 483–488.

Mullen, K. T., & Boulton, J. C. (1992b). Interactions between colour and luminance contrast in the perception of motion. Ophthalmic and Physiological Optics, 12(2), 201–205.

Mullen, K. T., Yoshizawa, T., & Baker Jr, C. L. (2003). Luminance mechanisms mediate the motion of red–green isoluminant gratings: The role of “temporal chromatic aberration”. Vision Research, 43(11), 1237–1249.

Nassi, J.J. & Callaway, E.M. (2009). Parallel processing strategies of the primate visual system. Nature Rev 360(10), 360–372.

Ogboso, Y. U., & Bedell, H. E. (1987). Magnitude of lateral chromatic aberration across the retina of the human eye. Journal of the Optical Society of America A, 4(8), 1666–1672.

Peichl, L. (1991). Alpha ganglion cells in mammalian retinae: common properties, species differences, and some comments on other ganglion cells. Visual neuroscience, 7(1-2), 155–169.

Pelli, D. G. (1997). The VideoToolbox software for visual psychophysics: transforming numbers into movies. Spatial Vision 10, 437–442 (1997).

Perry, V., Oehler, R., & Cowey, A. (1984). Retinal ganglion cells that project to the dorsal lateral geniculate nucleus in the macaque monkey. Neuroscience, 12(4), 1101–1123.

Prins, N., & Kingdom, F. A. (2018). Applying the model-comparison approach to test specific research hypotheses in psychophysical research using the Palamedes toolbox. Frontiers in psychology, 9, 1250.

Quigley, H. A., Dunkelberger, G. R., & Green, W. R. (1988). Chronic human glaucoma causing selectively greater loss of large optic nerve fibers. Ophthalmology, 95(3), 357–363.

Quigley, H. A., Sanchez, R. M., Dunkelberger, G. R., L’Hernault, N. L., & Baginski, T. A. (1987). Chronic glaucoma selectively damages large optic nerve fibers. Investigative ophthalmology & visual science, 28(6), 913–920.

Ramachandran, V.S. & Gregory, R.L. (1978). Does colour provide an input to human motion perception ? Nature 275: 55–56.

Ramirez, A. L., Thompson, L. W., Rosenberg, A., & Baker Jr, C. L. (2022). Behavioral signatures of Y-like neuronal responses in human vision. Scientific reports, 12(1), 19116.

Robson, J. G. (1966). Spatial and temporal contrast-sensitivity functions of the visual system. Journal of the optical society of America, 56(8), 1141–1142.

Rodieck, R. (1979). Visual pathways. Annual Review of neuroscience, 2(1), 193–225.

Rosenberg, A., Husson, T. R., & Issa, N. P. (2010). Subcortical representation of non-Fourier image features. Journal of Neuroscience, 30(6), 1985–1993.

Rosenberg, A., & Issa, N. P. (2011). The Y cell visual pathway implements a demodulating nonlinearity. Neuron, 71(2), 348–361.

Rosenberg, A., & Talebi, V. (2009). The primate retina contains distinct types of Y-like ganglion cells. Journal of Neuroscience, 29(16), 5048–5050.

Rynders, M. C., Navarro, R., & Losada, M. A. (1998). Objective measurement of the off-axis longitudinal chromatic aberration in the human eye. Vision Research, 38(4), 513–522.

Schwiegerling, J. (2000). Theoretical limits to visual performance. Survey of ophthalmology, 45(2), 139–146.

Shapley, R., & Perry, V. H. (1986). Cat and monkey retinal ganglion cells and their visual functional roles. Trends in Neurosciences, 9, 229–235.

Shooner, C., & Mullen, K. T. (2020). Enhanced luminance sensitivity on color and luminance pedestals: Threshold measurements and a model of parvocellular luminance processing. Journal of Vision, 20(6), 12.

Shou, T., Liu, J., Wang, W., Zhou, Y., & Zhao, K. (2003). Differential dendritic shrinkage of α and β retinal ganglion cells in cats with chronic glaucoma. Investigative ophthalmology & visual science, 44(7), 3005–3010.

Smith, W. J. (1990). Modern Optical Engineering (2 ed.). McGraw-Hill.

Stromeyer 3rd, C., Chaparro, A., Tolias, A., & Kronauer, R. (1997). Colour adaptation modifies the long-wave versus middle-wave cone weights and temporal phases in human luminance (but not red-green) mechanism. The Journal of physiology, 499(1), 227–254.

Stromeyer 3rd, C., Gowdy, P., Chaparro, A., Kladakis, S., Willen, J., & Kronauer, R. (2000). Colour adaptation modifies the temporal properties of the long-and middle-wave cone signals in the human luminance mechanism. The Journal of physiology, 526(Pt 1), 177–194.

Stromeyer 3rd, C., Kronauer, R., Ryu, A., Chaparro, A., & Eskew Jr, R. (1995). Contributions of human long-wave and middle-wave cones to motion detection. The Journal of physiology, 485(1), 221–243.

Thibos, L.N. (1987). Calculation of the influence of lateral chromatic aberration on image quality across the visual field. J. Opt. Soc. Am. A 4, 1673–1680.

Thibos, L. N., Bradley, A., & Zhang, X. (1991). Effect of ocular chromatic aberration on monocular visual performance. Optometry and Vision Science, 68(8), 599–607.

Thibos, L. N., Walsh, D., & Cheney, F. (1987). Vision beyond the resolution limit: aliasing in the periphery. Vision Research, 27(12), 2193–2197.

Tsujimura, S., Shioiri, Y. Hirai, Yaguchi, H.. (1999). Selective cone suppression by the L-M- and M-L-cone-opponent mechanisms in the luminance pathway. Journal of the Optical Society of America A, 16, 1217–1228.

Tsujimura, S., Shioiri, S., Hirai, Y., Yaguchi, H.. (2000). Technique to investigate the temporal phase shift between L- and M-cone inputs to the luminance mechanism. Journal of the Optical Society of America A, 17, 846–857.

Walsh, G. (1988). The effect of mydriasis on the pupillary centration of the human eye. Ophthalmic and Physiological Optics: the journal of the British College of Ophthalmic Opticians (Optometrists), 8(2), 178–182.

Watson, A. B., & Ahumada, A. J. (2016). The pyramid of visibility. Electronic Imaging, 28, 1–6.

Yoshizawa, T., Mullen, K. T., & Baker Jr, C. L. (2000). Absence of a chromatic linear motion mechanism in human vision. Vision Research, 40(15), 1993–2010.

Yoshizawa, T., Mullen, K. T., & Baker Jr, C. L. (2003). Failure of signed chromatic apparent motion with luminance masking. Vision Research, 43(7), 751–759.

Yücel, Y. H., Zhang, Q., Gupta, N., Kaufman, P. L., & Weinreb, R. N. (2000). Loss of neurons in magnocellular and parvocellular layers of the lateral geniculate nucleus in glaucoma. Archives of ophthalmology, 118(3), 378–384.

Zhang, P., Wen, W., Sun, X., & He, S. (2016). Selective reduction of fMRI responses to transient achromatic stimuli in the magnocellular layers of the LGN and the superficial layer of the SC of early glaucoma patients. Human brain mapping, 37(2), 558–569.

Zhou, Y.-X., & Baker Jr, C. L. (1994). Envelope-responsive neurons in areas 17 and 18 of cat. Journal of neurophysiology, 72(5), 2134–2150.

Zhou, Y.-X., & Baker Jr, C. L. (1996). Spatial properties of envelope-responsive cells in area 17 and 18 neurons of the cat. Journal of neurophysiology, 75(3), 1038–1050.

